# Gene- and pathway-based association tests for multiple traits with GWAS summary statistics

**DOI:** 10.1101/052068

**Authors:** Il-Youp Kwak, Wei Pan

## Abstract

To identify novel genetic variants associated with complex traits and to shed new insights on underlying biology, in addition to the most popular single SNP-single trait association analysis, it would be useful to explore multiple correlated (intermediate) traits at the gene-or pathway-level by mining existing single GWAS or meta-analyzed GWAS data. For this purpose, we present an adaptive gene-based test and a pathway-based test for association analysis of multiple traits with GWAS summary statistics. The proposed tests are adaptive at both the SNP-and trait-levels; that is, they account for possibly varying association patterns (e.g. signal sparsity levels) across SNPs and traits, thus maintaining high power across a wide range of situations. Furthermore, the proposed methods are general: they can be applied to mixed types of traits, and to Z-statistics or p-values as summary statistics obtained from either a single GWAS or a meta-analysis of multiple GWAS. Our numerical studies with simulated and real data demonstrated the promising performance of the proposed methods.

The methods are implemented in R package aSPU, freely and publicly available on CRAN at: https://cran.r-project.org/web/packages/aSPU/.

## 1 Introduction

In spite of the success of genome-wide association studies (GWAS) in identifying thousands of reproducible associations between single nucleotide polymorphism (SNPs) and complex diseases/traits, in general the identified genetic variants can explain only a small proportion of heritability (Manolio *et al*. 2009). A main reason is due to small effect sizes of genetic variants, raising both challenges and opportunities in developing more powerful analysis strategies. Among others, endeavors in the following three directions have been undertaken. First, due to polygenic effects (with small effect sizes) on complex traits, instead of the popular single SNP-single trait analysis, it may be more powerful to conduct gene-and pathway-level association tests (Lin and Tang, 2011; Wu *et al.* 2010; Pan *et al.* 2014; Li, *et al.* 2011; Gui *et al.* 2011; Li *et al.* 2012; Pan *et al.* 2015). However, most of the existing association tests are based on the use of individual-level genotypic and phenotypic data, while quite often only summary statistics for single SNPs are available. Thus, some association tests for a single trait but applicable to GWAS summary statistics have appeared, including GATES (Li *et al.* 2011), GATES-Simes (Gui *et al.* 2011), HYST (Li *et al.* 2012), and aSPUs and aSPUsPath (Kwak and Pan 2015). Second, while many GWAS have collected multliple (intermediate) traits, due to pleiotropic effects, multiple correlated (intermediate) traits, e.g. neuroimaging endophenotypes (Shen *et al.* 2010; Zhang *et al.* 2014), can be used to boost power and illuminate on underlying biological mechanisms as compared to popular disease-based single trait analyses; see a review by Yang and Wang (2013). Most of the existing association tests for multiple traits are based on individual-level data, (Basu *et al.* 2013; Tang and Ferreira 2012; Yang et al 2010; Zhang *et al.* 2014; Wang *et al.* 2015; Fan *et al.* 2015, 2016) with only few exceptions such as MGAS (Sluis *et al.* 2015) and metaCCA (Cichonska *et al.* 2016). Third, to increase the sample size, large consortia are being formed, aiming for meta analysis of multiple GWAS, for which often only summary statistics for single SNP-single trait associations, rather than individual-level genotypic and phenotypic data, are available. Hence it is necessary to develop methods that are applicable to only summary statistics. Motivated by the above three considerations, here we present such tests.

To our knowledge, there are only two existing tests that are for gene-or pathway-based analysis of multiple traits and applicable to summary statistics. MGAS (Sluis *et al.* 2015) uses an extended Simes procedure and behaves like a univariate minimum p-value approach, while metaCCA (Cichonska *et al.* 2016) is based on canonical correlation analysis (CCA) of multiple traits and multiple SNPs, which is related to MANOVA and GEE-score test (Zhang *et al.* 2014; Kim *et al.* 2016); the two tests may lose power in some situations with multiple but relatively sparse and weak association signals between the traits and SNPs (Pan *et al.* 2014; Zhang *et al.* 2014). Accordingly, it would be useful to extend adaptive tests for multiple trait-single SNP (Kim *et al.* 2015) or for single trait-multiple SNP associations (Kwak and Pan, 2015) with summary statistics, or for multiple trait-multiple SNP associations with individual-level data (Kim *et al.* 2016), to the current case of multiple trait-multiple SNP associations with only GWAS summary statistics, which is the aim here. In addition, we propose a novel Monte Carlo simulation method based on a matrix normal distribution to estimate the p-values for our proposed tests, which is well justified by known asymptotic theory that is suitable for large GWAS. In our proposed approach, we use a reference panel to estimate linkage disequilibrium (LD) among physically nearby SNPs; in contrast, metaCCA uses a similar method to estimate a joint covariance matrix for both the multiple traits and multiple SNPs, possibly explaining why it requires a large sample size of the reference panel to perform well, as to be confirmed in our later simulations. We also note that in MGAS, instead of individual-level genotypic data in a reference panel, p-values as summary statistics are used to empirically estimate LD among SNPs, which may lead to non-positive definite correlation matrices as numerically shown in Kwak and Pan (2016).

Finally we note that our proposed methods are general with a wide range of applications. For example, the multiple traits can be mixed types: some may be quantitative while others binary; the summary statistics for single SNP-single trait associations, as either Z-statistics or p-values, can be obtained from either a singe GWAS or a meta-analysis of multiple GWAS (with any valid test being applied). It is noteworthy to point out that the current version of metaCCA requires an equal sample size for all SNP-trait pairs, which is too restrictive for meta-analyzed GWAS. For example, the sample sizes for the SNP-trait summary statistics in a real dataset to be analyzed varied dramatically, rendering the non-applicability of metaCCA.

We will validate the proposed methods using the Welcome Trust Case Control Consortium (WTCCC) GWAS data (WTCCC 2007), then illustrate their applications to a meta-analyzed dataset from the Genetic Investigation of ANthropometric Traits (GIANT) consortium (Randall *et al.* 2013). We will compare our methods with MGAS and metaCCA, demonstrating the promising performance and advantages of our methods.

## 2 Methods

### 2.1 Notation

Suppose there are *d* SNPs (e.g. in a gene for gene-based testing) with additive genotype scores G = (*G*_1_, ⋯, *G*_*d*_)′, where *G*_*j*_ is the number of minor alleles of the *j*th SNP; there are *m* > 1 quantitative or binary phenotypes *Y* = (*Y*_1_, …,*Y*_*m*_)′; let *C* = (*C*_1_, ⋯, *C*_*l*_)′ denotes a set of covariates. We first consider one phenotype *Y*_*h*_ by applying a generalized linear model:

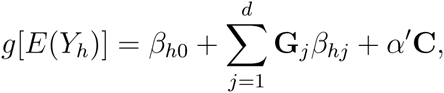

where *g*() is a canonical link function (i.e. the identity function for a quantitative trait, or a logit function for a binary trait). We are interested in testing *H*_0_: *β*_*hj*_ = 0 for all *h* = 1, ⋯, *m* and *j* = 1, ⋯, *d*).

For a given dataset {(*Y*_*ih*_, *G*_*i*_, *C*_*i*_): *i* = 1,…,*n*} with *n* subjects, the score vector **U**_**h**_ = (*U*_*h*1_, ⋯, *U*_*hd*_)′ for *β*_*h*_ is

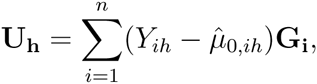

where 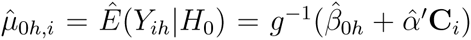 is the estimated mean of *Y*_*ih*_ in the null model (under *H*_0_).

Kim *et al.* (2016) constructed an adaptive test for multi-trait and multi-SNP association using the score vector. However, in the current context without individual-level data, we cannot directly calculate *U*_*hj*_’s as given in the formula.

Here we assume that we only have summary statistics, say an *m* × *d* matrix of Z scores, **Z**. Each element *Z*_*hj*_, from the ith row and *j*th column of **Z**, represents a Z score for testing association between the *h*th phenotype and the *j*th SNP. A Z score is (asymptotically) a weighted version of an element in the score vector: 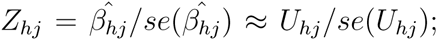 the approximation is based on the asymptotic equivalence between the Wald test and the Score test. Taking the Z scores in place of the score vector has been proposed to test for multitrait-single SNP associations (Kim *et al.* 2015) and single trait-multiple SNP associations (Kwak and Pan 2015).

### 2.2 Gene-based tests

We extend the gene-based tests based on individual-level data (Kim *et al.* 2016) to those based on summary statistics. Specifically, we define a test statistic for single trait-multiple SNP association and that for multiple trait-multiple SNP association as

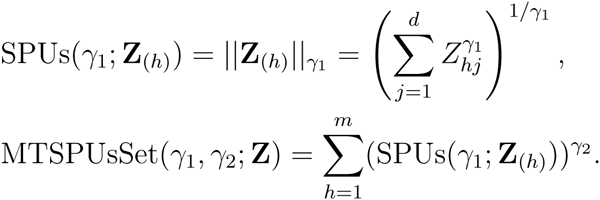

where **Z**_(*h*)_ represents the *h*th row vector of matrix **Z**; i.e. the Z scores for the *h*th trait. Two scalars *γ*_1_ ≥ 1 and *γ*_2_ ≥ 1 controls the extents of weighting on the SNPs and traits respectively. For example, a larger *γ*_1_ (or *γ*_2_) is expected to yield higher power if there are a smaller number of the SNPs (or traits) with truly non-zero associations (i.e. with the corresponding (*β*_*hj*_ ≠ 0). As discussed in more details in Kim *et al.* (2016), MTSPUsSet(1, 1) is like a burden test (Shen *et al.* 2010), while MTSPUsSet(*γ*_1_,*γ*_2_) for large values of *γ*_1_ and *γ*_2_ is effectively equivalent to a univariate minimum p-value test on all single SNP-single trait pairs; MTSPUsSet(2, 2) is closely related to a variance-component score test in kernel machine regression (Maity *et al.* 2012) and nonparametric MANOVA or distance-based regression (McArdle and Anderson 2001; Wessel and Schork 2006; Schaid 2005).

Since the optimal values of (*γ*_1_,*γ*_2_) are unknown, we propose an adaptive test to data-adaptively choose (*γ*_1_,*γ*_2_):

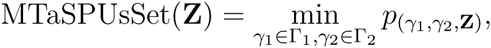

where *p*(*γ*_1_,*γ*_2_),**Z**) is the p-value for MTSPUsSet(*γ*_1_, *γ*_2_, **Z**), and by default we use Γ_1_ = {1, 2, 4, 8} and Γ_2_ = {1, 2, 4, 8}.

A main innovation here is to use a matrix normal distribution (Gupta and Nagar 1999; Zhou 2014) to obtain p-values based on the known asymptotic normal distribution of the Z scores under *H*_0_. Specifically, denote **Z**_(*i*)_ as the *i*th row vector, and **Z**_*j*_ as the *j*th column vector (i.e. the Z scores for *j*th SNP) of Z. If the sample size is large (with relatively small numbers of traits and SNPs), by the standard asymptotics for the Z scores, it is reasonable to assume that the null distribution of Z is a matrix normal distribution:

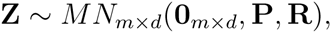

where 0_*m*×*d*_ is the *m* × *d* matrix with 0’s. It is equivalent to saying that

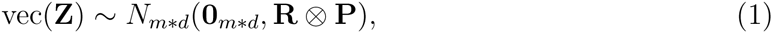

where vec(**Z**) is formed by stacking the columns of **Z**, ⊗ is the Kronecker product, and 0_*m*\**d*_ is a 0 vector of length *m* * *d*.

From equation (1), We see that 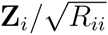 follows a normal distribution with mean 0 and covariance matrix **P**, and that 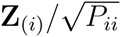 follows a normal distribution with mean 0 and covariance matrix **R** (Zhou, 2014). Since **P** and **R** are correlation matrices with *R*_*ii*_ = *P*_*ii*_ = 1, we obtain

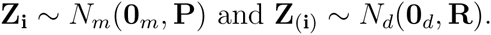

Following Kim *et al.* (2015), we propose excluding the SNPs with small p-values (e.g. < 0.05) and using a large subset of the remaining null SNPs to estimate **P** with the sample correlation matrix of the Z scores. For **R**, as shown by Kwak and Pan (2015) and others, it can be approximated by the sample correlation matrix of the SNPs using a reference panel similar to the study population. For example, we used 1000G Phase I version 3 Shapeit2 Reference data downloaded from the KGG software website (Li *et al.* 2012); it contains about 81.2 million polymorphic markers on 2,504 samples released in September 2014. By default, we used 379 CEU (Utah Residents with Northern and Western Ancestry) samples.

Finally we note that, based on the asymptotic null distribution of vec(**Z**) in (1), we can construct a score test (if **Z** is obtained by the univariate score test or its asymptotically equivalent tests like the Wald test):

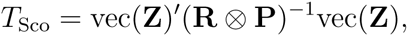

which has an asymptotic 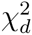 with degrees of freedom *d* = rank(**R** ⊗ **P**); if **R** ⊗ **P** is not of full rank, a generalized inverse is used in *T*_Sco_.

As discussed in Zhang *et al.* (2014) and Kim *et al.* (2016), the score test is similar to CCA and MANOVA, hence we expect that *T*_Sco_ will perform similarly to metaCCA, as to be confirmed. Furthermore, the score test behaves differently from the aSPU test; neither can dominate the other with higher power in all applications. Hence, it might be useful to combine the two tests as

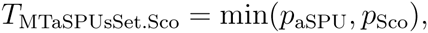

where *p*_aSPU_ and *p*_Sco_ are the p-values of the MTaSPUsSet and *T*_Sco_ respectively; as to be shown, the p-values of all the tests could be obtained simultaneously in a single layer of Monte Carlo simulations.

### 2.3 Pathway-based tests

We extend the pathway-based multi-trait association tests of Kim *et al.* (2016) to the case with only GWAS summary statistics. Given a pathway *S* with |*S*| genes, we partition the Z score matrix as **Z** = (**Z**′_(1)_, ⋯, **Z**′_(*m*)_)′ with **Z**_(*i*)_ as the *i*th row vector (i.e. Z scores for the *i*th trait). **Z**_(*i*)_ is further partitioned at the gene level to **Z**_(*i*)_ = (**Z**′_(*i*1)_, **Z**′_(*i*2)_, ⋯, **Z**′_(*i*|*S*|)_)′, and at the SNP level to (**Z**_(*ig*)_ = (*Z*_(*ig*)1_, *Z*_(*ig*)2_, ⋯, *Z*_(*ig*)*d*_*g*__) (for the *d*_*g*_ SNPs in gene *g*).

We define the gene-and pathway-based tests for a single trait and then for multiple traits as

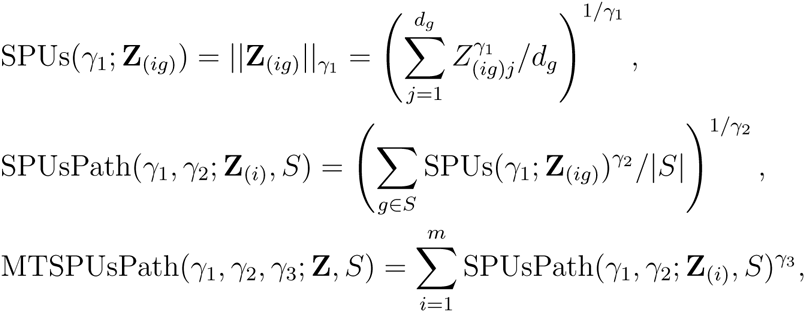

where the three integers *γ*_1_ ≥ 1, *γ*_2_ ≥ 1 and *γ*_3_ ≥ 1 are used to adaptively weight the SNPs, genes and traits respectively. For example, a larger *γ*_1_ (or *γ*_2_, or *γ*_3_) is more effective when there are a smaller number of truly associated SNPs (or genes, or traits).

To adaptively choose (*γ*_1_,*γ*_2_,*γ*_3_), we propose a pathway-based adaptive test as

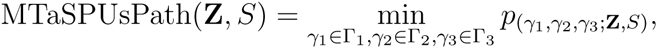

where p(*γ*_1_,*γ*_2_,*γ*_3_;**Z**,*S*) is the p-value of MTSPUsPath(*γ*_1_,*γ*_2_,*γ*_3_; **Z**, *S*), and by default we use Γ_1_ = {1,2, 4,8}, Γ_2_ = {1,2, 4, 8} and Γ_3_ = {1, 2, 4, 8}.

### 2.4 P-value calculations

Monte Carlo simulations are used to obtain the p-values for all the tests, including MTaSPUsSet or MTaSPUsSetPath, in a single layer of simulations. Briefly, after estimating **P** and **R**, first we simulate null scores **Z**^(*b*)^ ~ *M N*_*m*×*d*_ (0_*m*×*d*_, **P**, **R**) for *b* = 1, ⋯, *B*. Then we use the null scores to calculate the null test statistics, from which the p-values can be calculated (Kwak and Pan 2016). A larger *B* is needed to estimate a smaller p-value.

We generate a matrix normal variate **Z**^(*b*)^ in the following way (Zhou 2014). We first generate an *n* × *d* matrix **L** with each element independently from a standard univariate normal distribution with mean 0 and variance 1; that is,. **L** ~ *M N*_*m*×*d*_ (0_*m*×*d*_,*I*_*m*_, *I*_*d*_). Then we obtain **Z**^(*b*)^ = **DLE′**, where **D** and **E** are Cholesky decompositions of **P** and **R** with **P** = **DD′** and **R** = **EE′**.

Specifically, for MTaSPUsSet,

- Step 1.Generate independent **Z**^(*b*)^ ~ *M N*_*m*×*d*_ (0_*m*×*d*_, **P**, **R**) for *b* = 1, ⋯, *B*;
- Step 2.Calculate the null test statistics MTSPUsSet(*γ*_1_,*γ*_2_, **Z**^(*b*)^);
- Step 3.The p-value for MTSPUsSet(*γ*_1_,*γ*_2_; **Z**) is

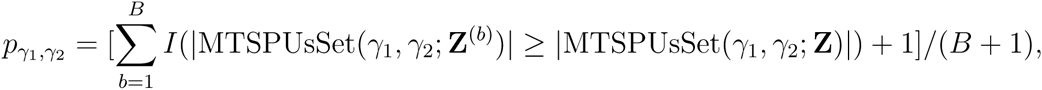

and that for MTSPUsSet(*γ*_1_,*γ*_2_; **Z**^(*b*)^) is

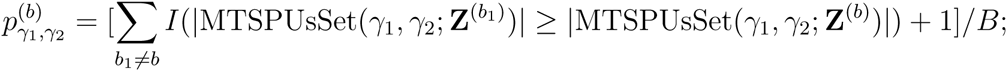
- Step 4.Calculate the null and observed test statistics

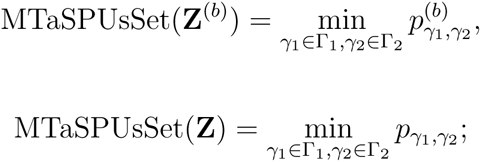
- Step 5.Finally the p-value for the MTaSPUsSet test is

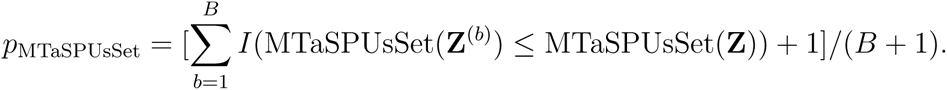

A similar procedure is used to obtain the p-values for MTSPUsPath and MTaSPUsPath. When only p-values for single SNP-single trait associations, instead of Z statistics, are available as summary statistics, we use |*Z*| = Φ^−1^(1 – *P*/2), where Φ is the cumulative distribution function of the standard univariate normal distribution; we replace all *Z*’s with |*Z*|’s to calculate the test statistics.

## 3 Results

### 3.1 Simulations

To demonstrate the validity and performance of our proposed methods, we designed a “Control-Control” experiment using the Welcome Trust Case Control Consortium (WTCCC) GWAS data for Crohn’s disease (CD) (Consortium 2007; Kwak and Pan 2016). The WTCCC GWAS dataset contains about 3,000 controls with a total of 500,568 SNPs. Following the WTCCC’s quality control (QC) recommendations, we removed subjects and SNPs that did not pass the QC criteria, resulting in 469,612 SNPs in 2,938 control subjects. We further removed SNPs with MAF<5% since we had only 379 samples in our reference panel to infer the LD structure for a set of SNPs. We considered 4,572 unique genes in 186 KEGG pathways to check type 1 error rates of our gene-based test. A total of 64,557 SNPs were mapped to these genes.

We simulated multiple traits using a multivariate normal distribution with mean 0 and correlation matrix in Equation (3) of Figure S1, which was estimated based on the GIANT data for women. We generated a set of six traits for each of the 2938 control subjects. Then we calculated the univariate Z scores for all 64,557-6 SNP-trait pairs. A Monte Carlo simulation size of *B* = 10^5^ was used to calculate the p-values.

For each gene (or pathway), **R** was estimated from the 1000 Genome Project CEU samples. To estimate **P**, we excluded the SNPs with p-values < 0.05 and used the remaining 48,669 SNPs. Equation (1) of Figure S1 is the estimate for **P**. This estimate is close to the true value shown in Equation (3) of Figure S1, **P**_**w**_. We pruned SNPs in high LD by removing any SNP if it was correlated with another SNP with an absolute value of Pearson’s correlation coefficient larger than 0.95.

### 3.2 Gene-based tests

We first investigated the effects of the choice of the reference panel on estimating LD among SNPs, i.e. **R** for each gene. We considered three scenarios: 1) using the whole 2938 WTCCC controls as the reference panel as an ideal case; 2) using only a random set of 100 WTCCC control samples as the reference panel to see whether a sample size as low as 100, close to that of many published reference panels, was sufficient to obtain accurate estimates; 3) using the 1000 Genomes Project CEU samples with 379 individuals as the reference panel, a more realistic scenario without individual level data.

Figure S2 shows the QQ plots of the p-values of the MTaSPUsSet test based on each of the three ways to estimate the SNP correlation matrix. We can see that all three plots looked reasonable with the estimated inflation factor λ’s as 1.01, 0.99 and 0.99 respectively, all close to 1. It was confirmed that the type I error rates seemed to be well controlled in all cases.

Next we further compared the results as shown in Figure 4. By comparing the results between using the WTCCC whole control samples and using only 100 samples as reference panel, we conclude that taking only 100 samples from the original whole dataset seemed to perform well; the Pearson correlation (*r*) between the two was 0.99. The top right and bottom left panels compare the results between using the WTCCC whole data, WTCCC 100 samples and 1000 Genome Project CEU samples as the reference panel; again they showed high degrees of mutual agreement with a Pearson correlation coefficient as high as 0.97 and 0.98 respectively. In the bottom right panel, we further compared the results of MTaSPUsSet with only summary statistics (using the 1000 Genome Project CEU samples as the reference panel) to a similar GEE-based adaptive test with individual-level data (Kim *et al.* 2016). Although the agreement was reasonably high with a Pearson correlation coefficient of 0.9, there were some differences, indicating that cautions are needed when using summary statistics.

We also tried metaCCA (Cichonska *et al.* 2016) and TSco on the simulated data, and found that both might not work well when the sample size of the reference panel was small. We used 1) the whole 2938 WTCCC controls as an ideal case; 2) 100-2000 samples from the WTCCC control data; 3) using the 1000 Genome Project CEU samples, respectively, as the reference panel. We used “metaCcaGp” function in the R version of metaCCA at: https://bioconductor.org/packages/devel/bioc/html/metaCCA.html. Figures S3 and S4 show the QQ plots for each scenario. In particular, it showed that even a sample size of 500 drawn from the WTCCC control data or of 379 for the 1000 Genome Project CEU samples might not be large enough; because of this reason, we would not apply the tests (and thus MTaSPUsSet.Sco either) to the real data.

Importantly, it was confirmed that metaCCA and *T*_Sco_ gave almost the same p-values, as shown in Figure S5

### 3.3 Pathway-based tests

For evaluations, we designed a control-control experiment using the WTCCC CD data. We randomly chosen 3 to 15 genes from the WTCCC data to form a pathway. We applied the MTaSPUsPath test to each of 319 pathways. Simulations were conducted with different reference panels used to estimate **R**, similar to what was done for gene-based testing.

Figure S6 compares the results of MTaSPUsPath with various reference panels, and of a similar pathway-based adaptive test called GEE-aSPUpath based on individual-level data (Kim *et al.* 2016). Similar conclusions to those for the gene-based MTaSPUsSet test can be drawn.

### 3.4 Analysis of GIANT data

We applied the MTaSPUsSet test to the summary statistics for sex stratified anthropometrics data from The Genetic Investigation of ANthropometric Traits (GIANT) consortium (Randall *et al.* 2013). The data contain the p-values of 2.7 million SNPs with each of six anthropometric traits that are well established to represent the body size and shape: height, weight, BMI, waist circumference (WC), hip circumference (HIP), and waist-hip circumference ratio (WHR).

The original study was based on a single SNP–single trait association analysis (Randall *et al.* 2013). Instead, we applied two gene-based association tests on the six traits (height, weight, BMI, WC, HIP and WHR) for men and for women separately: our proposed MTaSPUsSet and MGAS of Sluis *et al.* (2015). Since all study participants were of European ancestry, we used the 1000 Genome Project CEU samples as the reference panel for both methods.

First, for MTaSPUsSet, in total 2,722,976 SNPs were mapped to 17,562 genes (plus 2-kb upstream and 2-kb downstream regions). We set the genome-wide significance threshold at 0.05/17562 = 2.85 × 10^−6^ based on the Bonferroni correction. We pruned SNPs in high LD by removing any SNP if it was correlated with another SNP with an absolute value of Pearson’s correlation coefficient larger than 0.95. For each gene, the correlations among the SNPs, **R**, were estimated from the 1000 Genome Project CEU samples. The correlations among the six traits were estimated based on 1,454,615 null SNPs with non-significant Z scores for men and women respectively as shown in Figure S1.

A stage-wise simulation strategy was used to calculate the p-values for each gene. We started with the simulation number *B* = 10^4^; we sequentially increased *B* to 10^5^, then 10^6^ and finally 10^7^ if a gene’s p-value was less than 0.003, 0.0003 and 0.00003 respectively.

The MTaSPUsSet test identified a total of 137 genes to be genome-wide significant for men or women: 81 for men, 125 for women and 69 for both. As a comparison, for single SNP–single trait analysis, we used a genome-wide significance threshold of 5 × 10^−8^/6 based on a Bonferroni adjustment for six traits, yielding in total 1298 significant SNPs (with 623 SNPs mapped to 62 genes) for men, and 2072 significant SNPs (with 990 SNPs mapped to 97 genes) for women. Although there were many common genes (i.e. 53 and 85 for men and women) identified by both methods, the proposed MTaSPUsSet test identified more genes (Table S1). In particular, to demonstrate the sex differences of genetic effects, the new test pinpointed 12 and 56 significant genes uniquely and specifically for men and women respectively; in contrast, the popular and standard single SNP–single trait analysis identified 20 and 55 genes uniquely for men and women respectively. The smaller number of men-specific genes identified by the new test could be due to its higher power: it is reasonable to assume that some of the identified sex-specific genes are false positives due to inadequate power for either sex, though further validations are needed.

Next, we applied MGAS of Sluis *et al.* (2015) using “kgg” software. The same 2-kb upstream and 2-kb downstream regions were used in mapping the SNPs to each gene, and the same estimated trait correlation matrices were used. However, for unknown reasons, only in total 969,832 SNPs were mapped to 6,424 genes, compared to ours of mapping 2,722,976 SNPs to 17,562 genes. Accordingly, the genome-wide significance threshold was set at 0.05/6424 = 7.78 x 10^−6^ based on the Bonferroni correction. In total only 19 genes were identified by MGAS to be significant: 16 genes for women and 8 for men.

For a fair comparison between MTaSPUsSet and MGAS, we examined more closely the 17,562 and 6,424 mapped genes for each method. There were 5197 shared genes commonly mapped by both methods; many of the 6,424 “kgg” genes starting with “LOC” and “LINC” were not in the MTaSPUsSet set of the 17,562 genes. We decided to apply both methods to the common set of the 5197 genes. The genome-wide significance level was set at 0.05/5197 by the Bonferroni adjustment.

Figure 5 shows the Manhattan plots for men and women based on MGAS and MTaSPUsSet respectively. Although there were some shared and general patterns between the results of the two methods, MTaSPUsSet identified a larger number of significant genes: a total of 49 genes with 27 and 39 for men and women respectively. In contrast, MGAS identified only a total of 17 genes with 7 and 14 for men and women respectively. It might suggest that MTaSPUsSet was more powerful, though further validations are needed.

To further contrast the differences between the two tests, Table S2 lists the 17 significant genes identified by MGAS with the corresponding p-values from the two tests. Genes *LCORL*, *VTA1*, *BICD2*, *RASA2*, *GNA12*, *NCOA1*, *TNS1*, *CEP112*, *DNM3* and *RFWD2* were significant for women by both MGAS and MTaSPUsSet, and *LCORL*, *RASA2* and *NDUFS3* were significant for men by both tests, while *LCORL* and *RASA2* were significant for both men and women by both tests. Gene *LCORL* was known to be associated with anthropometric traits, including body height in African Americans (Carty *et al.* 2012), birth weight and adult height (Horikoshi *et al.* 2013); it is also a candidate gene for body weight in sheep (Al-Mamun *et al.* 2015) and body size in horse (Metzger *et al.* 2013).

Figure 6 shows the p-values of the univariate test on single trait-single SNP associations for some genes identified by MTaSPUsSet, along with the (*γ*_1_,*γ*_2_) values for the most significant MTSPUsSet(*γ*_1_,*γ*_2_) test for each gene. It can be seen that for genes *RPGRIP1L* and *RPS10-NUDT3*, since there were many moderately significant univariate p-values (for univariate trait-SNP associations) with a dense association pattern, small values (*γ*_1_, *γ*_2_) = (1, 2) or (2, 1) gave the most significant results. In contrast, for gene *DNM3* with a larger number of SNPs, the association pattern was more sparse with main associations between some SNPs and trait height, larger values of (*γ*_1_, *γ*_2_) = (4, 8) or (8, 8) gave the most significant result. On the other hand, for gene *ZCCHC2*, due to the two or three highly significant univariate p-values between one or two SNPs and two traits, weight and BMI, any value of (*γ*_1_,*γ*_2_) would detect the overall association.

## 4 Discussion

We have presented new gene-and pathway-based adaptive association tests for multiple traits using only GWAS summary statistics. Our control-control experiments using the WTCCC genotype data with simulated multiple traits demonstrated that the type I error rates were well controlled. For the estimation of LD among SNPs (i.e. correlation matrix **R**), the choice of a reference panel (with individual-level genotypic data) would be a key for the performance. In the WTCCC control-control experiments, we compared three reference panels based on either the whole or a small subset of the original WTCCC control data, and the 1000 Genome Project CEU samples (with 379 subjects). The p-values calculated from the three reference panels were in general similar, but not exactly the same; the Pearson correlation coefficient of the log(p-values) between any two reference panels was at least 0.97, confirming that either the 1000 Genome Project CEU samples or a small subset of the control samples from the original population were sufficient for the WTCCC subject population.

We applied our gene-based MTaSPUsSet test to the meta-analyzed GIANT data. Since the participants in the GIANT data were of European and European American descent, the use of the 1000 Genome Project CEU panel was expected to be reasonable. The MTaSPUsSet test identified a total of 137 significant genes: 81 for men, 125 for women and 69 for both. As a comparison, for single SNP–single trait analysis identified 117 genes: 62 for men, 97 for women and 42 for both. MTaSPUsSet identified more genes. For more comparison, we also applied MGAS (Sluis *et al.* 2015) using the same reference panel, identifying only 19 significant genes using “kgg” software with a smaller set of the genes being mapped. For a fair comparison, we applied both MTaSPUsSet and MGAS to a common set of 5197 genes. MTaSPUsSet identified 27 and 39 significant genes for men and women respectively, compared to only 7 and 14 genes by MGAS, suggesting possible power gains by MTaSPUsSet. We also note that the other method metaCCA could not be applied to the GIANT data because it required a common sample size for all SNP-trait pairs, while the sample size for some SNPs ranged from around 200 to about 70, 000 across the traits.

## 5 Software

The proposed methods are implemented in R package aSPU, which is unique with many functions for association testing on a single trait or multiple traits versus a single SNP or a gene or a pathway, based on either individual-level data or GWAS summary statistics. It is available at https://github.com/ikwak2/aSPU. A python version is also available at https://github.com/ikwak2/aSPU_py.

## Acknowledgment

### Funding

This research was supported by NIH grants R01GM113250, R01HL105397 and R01HL116720, and by the Minnesota Supercomputing Institute.

---

**Figure 1:**
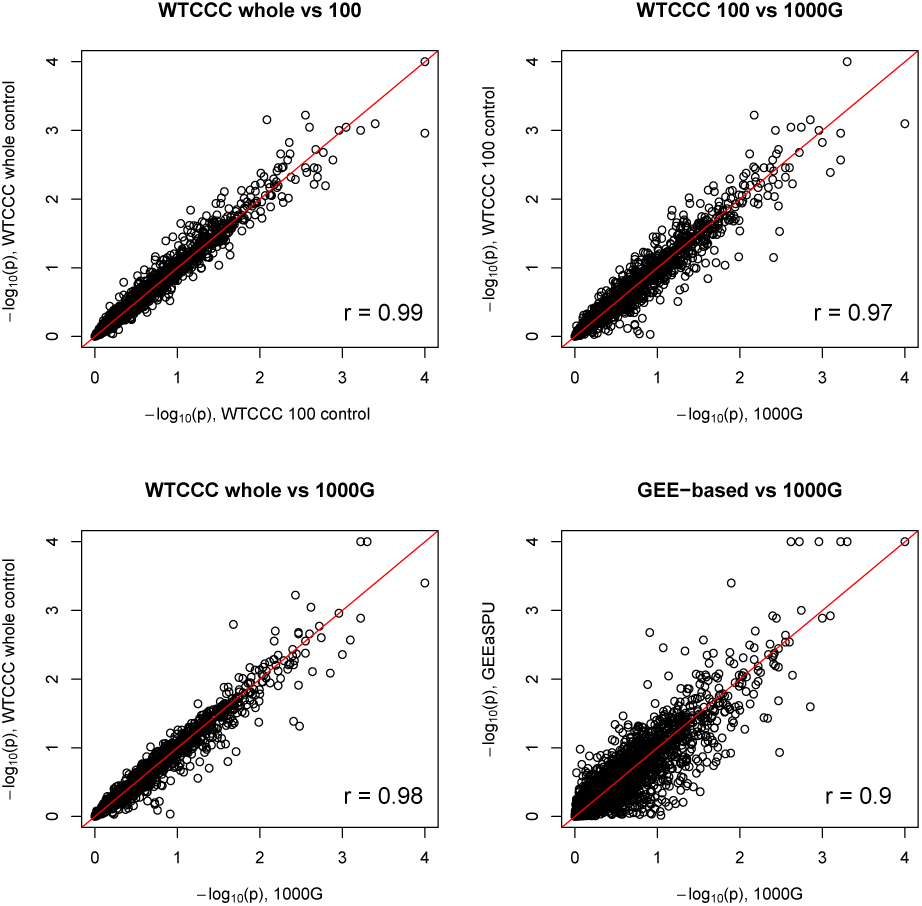
Comparison of the (log-transformed) p-values of MTaSPUsSet using various reference panels and that of the GEE-aSPU test using individual-level data.

**Figure 2:**
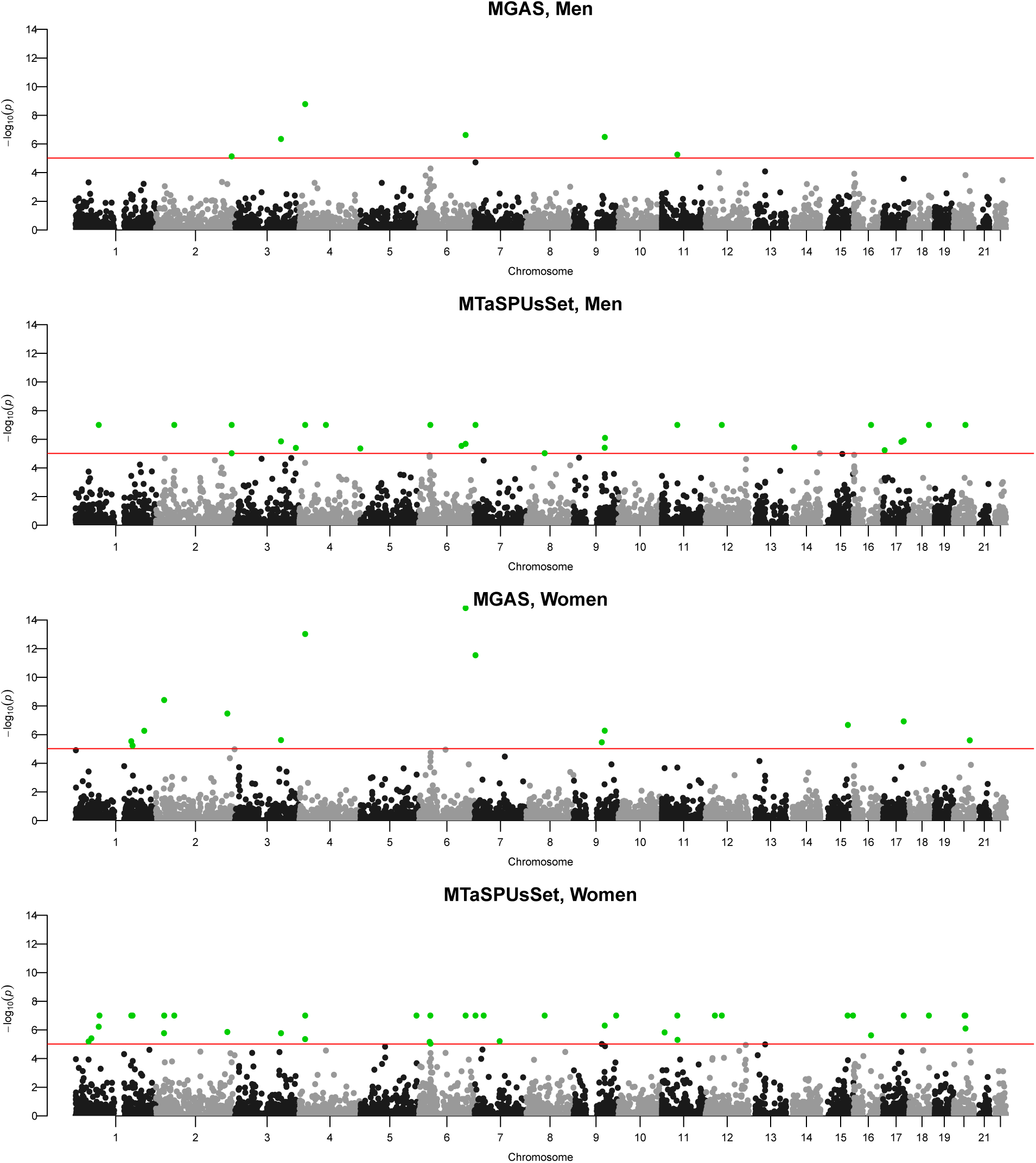
Manhattan plots for the GIANT data using MGAS and MTaSPUsSet on 5197 genes for men and women respectively.

**Figure 3:**
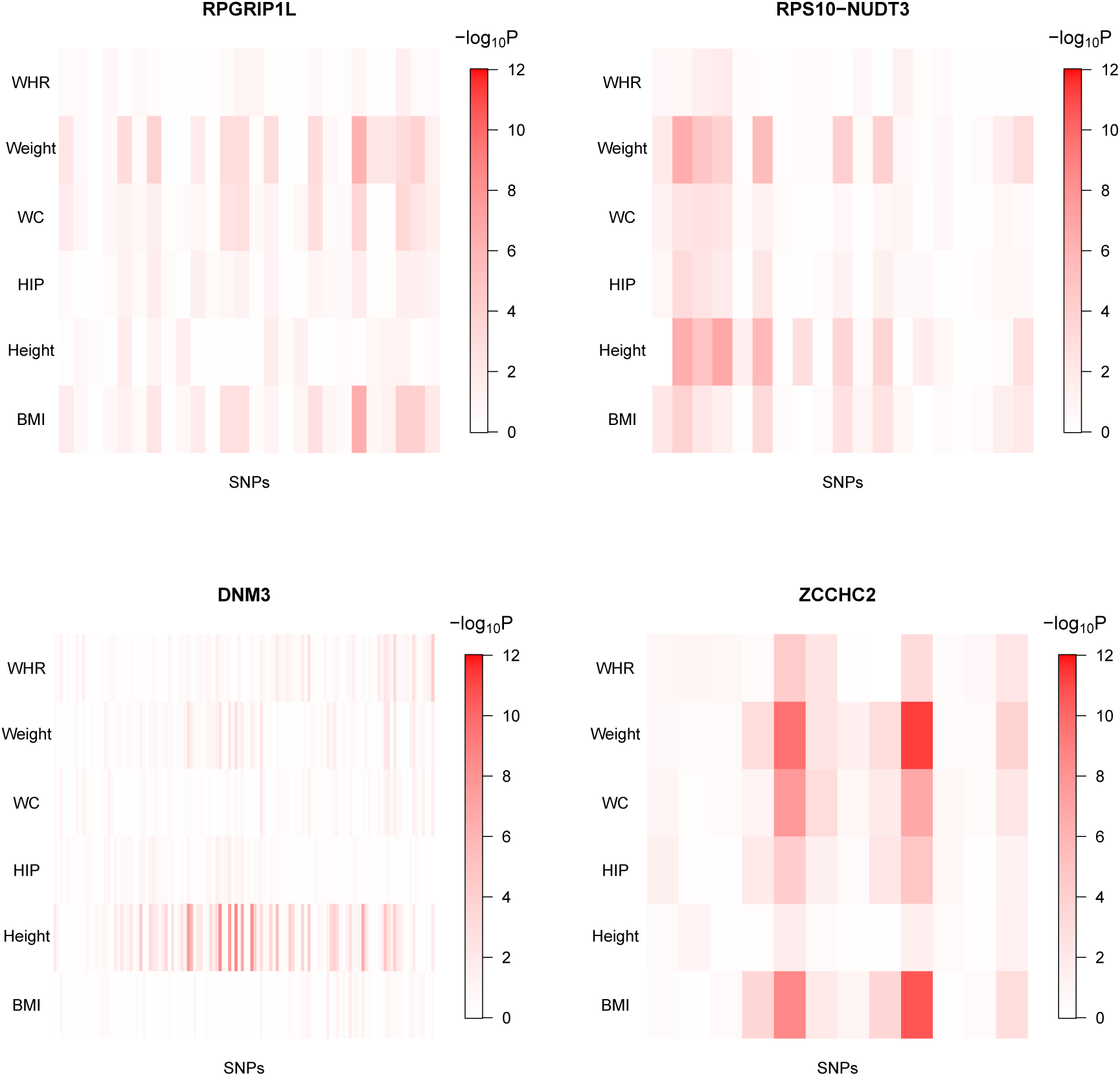
Log-transformed p-values of univariate SNP-trait associations for some genes identified by MTaSPUsSet. The most significant MTSPUsSet(γ_1_, γ_2_) test with the corresponding (γ_1_, γ_2_) values were (2, 1) for gene RPGRIP1L, (1, 2) for RPS10-NUDT3, (4, 8) or (8, 8) for *DNM3* and any value for *ZCCHC2*.

